# Spectral Phasor Analysis for Hyperspectral Imaging to elucidate Acridine Orange Photophysics in Cells

**DOI:** 10.1101/2023.07.11.548465

**Authors:** Marcela Díaz, Flavio R. Zolessi, Leonel Malacrida

## Abstract

Acridine Orange (AO) exhibits complex photophysical properties that have been utilized to study its interactions with RNA and DNA. Traditionally, AO has been used to stain cells, distinguishing nucleus and cytoplasm based on differences in DNA and RNA composition. Its labeling behavior varies depending on factors such as concentration and pH. AO fluorescence is rich and complex; therefore, acquiring the full spectrum or the fluorescence lifetime is beneficial for understanding its intricate photophysics. Nevertheless, handling 4D datasets can be challenging and often relies on sophisticated or highly parameterized models for their analysis. Model-free approaches, such as spectral phasor (SP) analysis for hyperspectral imaging (HSI), are well suited to navigating these analytical challenges. Using SP’s principles of linear combination and reciprocity, we identified spectral fingerprints of RNA and DNA in *in vitro* experiments that remain consistent in both live and fixed cells. Interestingly, SP analysis of AO fluorescence in live cells revealed a third spectral component at 640 nm. The linear properties of SP analysis enabled the quantification of AO bound to RNA, DNA, and the third component. Co-labeling with AO and LysoTracker suggested that some of these subcellular structures are associated with acidic compartments. Moreover, SP analysis distinguished AO self-interactions from its RNA binding in cytoplasmic compartments. This study establishes SP analysis as a robust tool for hyperspectral imaging of AO, thereby enhancing our understanding of its intricate photophysics and unlocking new opportunities for advanced *in vivo* cell imaging applications.

## 1. Introduction

Acridine Orange (AO) is a cationic dye with unique photophysical properties, making it a versatile tool for studying biological systems. Its spectral characteristics are highly dependent on parameters such as concentration, charge, and the surrounding molecular environment, as widely described in the literature (Kasten,1999). AO exists in two primary forms in aqueous solutions: at low concentrations (below 10⁻⁵ M), it predominantly forms monomers, with a maximum absorption peak at 490–494 nm and emission at 530 nm. At higher concentrations (above 10⁻² M), AO forms dimers with an absorption maximum at 465–467 nm, exhibiting red fluorescence with an emission peak at 640 nm (Markarian & Shahinyan, 2015; Shenderovich, 2019; Zelenin, 1999).

Since its first application in the 1940s to label cell nuclei in living and dead plant cells, AO fluorescence has been a cornerstone for studying nucleic acids. Metachromatic shifts observed with AO are attributed to its differential interactions with single and double stranded nucleic acids (Kasten,1999; Kapuscinski et al., 1982; Pierzyńska-Mach et al., 2014; Zelenin, 1999). When AO binds to double-stranded (ds) DNA or RNA at low dye-to-nucleic acid ratios, monomeric AO intercalates between base pairs, forming A-type complexes. These complexes are characterized by absorption at 503 nm and green fluorescence with a peak at 530 nm. Conversely, AO forms B-type complexes with single-stranded (ss) RNA or denatured DNA, characterized by absorption at 475 nm and red fluorescence with a peak at 640 nm. In this case, "stacking" interactions occur as AO binds to the phosphate backbone, leading to polymeric aggregates of dimers and multimers (Delic et al., 1991; Kasten, 1999; MacInnes & Uretz, 1966). At very high concentrations, RNA binding sites become saturated, leading to the precipitation of solid aggregates emitting red fluorescence (Kapuscinski et al., 1982; Plemel et al., 2017).

Under controlled pH and concentration conditions, AO is an effective cytochemical probe for nucleic acid labeling in fixed cells. In these cases, the nucleus (DNA) emits green fluorescence, while the cytoplasm and nucleoli (RNA) emit red fluorescence (Zelenin, 1999). However, labeling patterns differ in living cells when using low AO concentrations (5 × 10⁻⁶ M). In live cells, a bright green nucleus and nucleoli are observed, with prominent red cytoplasmic granules (Zelenin, 1999). First reported in 1944, these granules were initially attributed to AO–RNA complexes. Later studies in the 1960s identified them as lysosomes and other acidic compartments (Delic et al., 1991; Pierzyńska-Mach et al., 2014; Thomé et al., 2016; Zelenin, 1999). AO accumulates in acidic vesicles not solely due to low pH but also because of energy-dependent processes that trap protonated AO molecules within these compartments, resulting in red metachromatic shifts as the dye forms dimers and excimers (Hubenko et al., 2018; Pierzyńska-Mach et al., 2014; Thomé et al., 2016). Plemel et al. (Plemel et al., 2017) using hyperspectral imaging to demonstrate distinct spectral characteristics of AO in MO3 cells, with nuclear fluorescence peaking at 530 nm and cytoplasmic fluorescence peaking at 650 nm. To date, this remains one of the few studies leveraging hyperspectral imaging to explore AO fluorescence in live cells (Plemel et al., 2017).

Hyperspectral microscopy is an advanced imaging modality that combines the spatial resolution of optical confocal microscopy with the ability to collect detailed spectral information at each pixel of an image (Banu et al., 2024; Mangotra et al., 2023). This technique enables discrimination of overlapping fluorophores, identification of chemical compositions, and mapping of subtle spectral variations across complex biological samples (An et al., 2022; Dung et al., 2023; Mukhtar et al., 2025). Despite its potential, hyperspectral imaging presents technical challenges, including the need for specialized hardware and the processing of large, multidimensional datasets (Banu et al., 2024; Rodarmel & Shan, 2002). Traditional analysis methods such as Principal Component Analysis (PCA) and linear unmixing are commonly used but have limitations related to model assumptions, sensitivity to noise, and loss of information (Banu et al., 2024; Chang, 2003; Keshava & Mustard, 2002; Rodarmel & Shan, 2002).

The Spectral Phasor (SP) approach provides a model-free, intuitive method for analyzing hyperspectral data (Fereidouni et al., 2012; Malacrida & Gratton, 2018). By transforming the emission spectra into a phasor space based on Fourier components, SP enables direct visualization and comparison of spectral features without requiring prior knowledge of the sample (Torrado et al., 2022). The principles of linear combination and reciprocity inherent to the SP approach make it particularly well-suited for complex biological environments, where multiple fluorescent species may coexist and interact as in live-cell experiments (Castro-Castillo et al., 2020; Díaz & Malacrida, 2023; Lepanto et al., 2021; Malacrida et al., 2016; Malacrida & Gratton, 2018; Otaiza-González et al., 2022; Sena et al., 2017; Vorontsova et al., 2022).

In the present study, we applied hyperspectral confocal microscopy combined with spectral phasor analysis to investigate the metachromatic properties of AO in both fixed and live mammalian cells. This approach allowed us to spatially and spectrally resolve AO’s interactions with DNA, RNA, and intracellular compartments. By leveraging the capabilities of SP analysis, we were able to distinguish AO’s distinct molecular environments and provide new insights into the origin and distribution of its red-shifted fluorescence. Our findings demonstrate the utility of combining hyperspectral imaging with SP analysis for probing nucleic acid dynamics and subcellular organization in living cells, expanding the toolkit available for quantitative cytochemical investigations.

## 1. Materials and methods

### 1.1. Dyes and Reagents

Acridine Orange hemi (zinc chloride) salt (Sigma-Aldrich, #A6014) was prepared as a 27 mM stock solution in DMSO and used at a working concentration of 50 µM. For DNA and RNA experiments, the dye was diluted in phosphate-buffered saline (PBS), whereas for live-cell assays it was diluted in cell culture medium. LysoTracker™ Deep Red (ThermoFisher, #L12492) was employed for *in vitro* staining at a final concentration of 0.3 µM in cell culture medium. Fixation was performed using 4% paraformaldehyde in PBS (Sigma-Aldrich, #158127), and permeabilization was carried out with 0.1% Triton™ X-100 in PBS (Sigma-Aldrich, #X100). For nucleic acid reference samples, UltraPure Salmon Sperm DNA (ThermoFisher, #15632011) was used at concentrations of 0.1 and 10 mg/mL, while yeast RNA (Sigma-Aldrich, #10109223001) was diluted in PBS at concentrations of 0.7 and 2 mg/mL.

### 1.2. AO Binding to DNA or RNA

Acridine Orange (AO) stock solution was mixed with ultra-pure DNA or RNA solutions derived from yeast, both diluted in phosphate-buffered saline (PBS), to achieve a final AO concentration of 50 µM. The optimal concentration at which AO was fully bound to nucleic acids was determined by testing various concentrations of AO. The binding efficiency was evaluated by assessing the signal-to-noise ratio of AO and by monitoring shifts in the spectral phasor plot, specifically toward the blue, which indicated an increase in AO binding to DNA or RNA.

### 1.3. Cell culturing

NIH-3T3 cells (ATCC #CRL-1658™) were cultured in DMEM with GlutaMAX (ThermoFisher, #10566016), supplemented with 10% Donor Bovine Serum (ThermoFisher, #16030-074), 10 mM HEPES (ThermoFisher, #15630080), and 100 U/mL penicillin-streptomycin (ThermoFisher, #15140122). Cells were maintained at 37°C in a 5% CO2 atmosphere. For subculturing, cells were detached using Trypsin containing 0.25% EDTA (ThermoFisher, #25300054).

For imaging, cells were seeded onto 35 mm glass-bottom Petri dishes (Greiner Bio-One, #627870) at a density of 2.5 × 10⁵ cells per dish, 24 hours prior to imaging.

### 1.4. Cell staining

#### 1.4.1. NIH-3T3 live cells

AO staining was performed by incubating cells with the indicated AO concentration, diluted in cell culture medium, for 10 minutes at 37°C. After incubation, the cells were washed with fresh medium.

For LysoTracker™ Deep Red staining, cells were initially incubated with the dye for 10 minutes at 37°C. Following a wash with cell culture medium, confocal images were acquired. Subsequently, AO was added to the medium, and after another wash with fresh medium, additional confocal images were acquired.

#### 1.4.2. Fixed NIH-3T3 cells

Cells were fixed in 4% paraformaldehyde (PFA) for 10 minutes and then washed twice with PBS containing 0.1% Triton X-100. For staining, cells were incubated with the specified concentration of AO diluted in PBS (pH of the staining solution 7.4) for 10 minutes before imaging under the microscope.

### 1.5. Confocal microscopy and hyperspectral imaging

All spectral images were acquired using a Zeiss LSM 880 confocal microscope, equipped with a 63x oil immersion lens (NA 1.4, Plan Apochromat, DIC M27, Zeiss) and operated with the lambda mode of Zen Black v. 2.3 software (Carl Zeiss Microscopy GmbH, Jena, Germany). For AO excitation, a 488 nm Argon laser was used for and a 633 nm diode laser for LysoTracker™ Deep Red. A multiband pass main beam splitter (MBS 488/561/633) was employed. Emission spectra were captured sequentially using a spectral detector (GaAsP-PMT), with 22 consecutive steps (channels). Spectral images of 256 × 256 pixels were acquired at 10 nm intervals, covering the spectral range from 503 to 723 nm.

For AO-nucleic acid solutions, a 30 µL drop was placed on a coverslip and imaged using spectral confocal microscopy. The excitation was set to 488 nm (6.03 µW of laser energy delivered to the sample), utilizing the MBS 488. Spectral images (256 × 256 pixels) were acquired at 10 nm resolution, covering the 503 to 723 nm range. The pixel dwell time was 0.67 µs/pixel, and a line average of 2 was applied. For cell imaging with emission detection in two channels, a 488 nm Argon laser was used for excitation. Green fluorescence was detected in the range of 500 to 582 nm, and red fluorescence in the range of 583 to 630 nm, with the MBS 488. When evaluating LysoTracker™ Deep Red, excitation was performed using a 633 nm diode laser. A third emission channel was acquired in the range of 698 to 740 nm, using the MBS 488/561/633. Frame sizes of 512 × 512 pixels were acquired, pixel size of 50 nm, covering the spectral range from 503 to 723 nm (with 10 nm resolution).

During live-cell imaging, controlled environmental conditions were maintained at 37°C and 5% CO_2_. All images for fixed and live cells were captured with a pixel size between 130 and 100 nm, and the pixel dwell time was set to 0.85 μs/pixel. For live-cell experiments, the laser power delivered to the sample was 2.03 µW for the 488 nm laser and 3.87 µW for the 633 nm laser.

#### 1.5.1. Assessment of Autofluorescence in Unstained Cells

Spectral imaging of both fixed and live unstained cells was performed to verify that, at the laser power and detector gain levels used throughout this study, no significant autofluorescence signal was observed, either in the emission spectra or in the associated phasor plots.

### 1.6. Image processing

For images shown in the figures of this paper: Brightness and contrast adjustments, application of a smooth filter, summation of lambda channels for generating stack images, and placement of the scale bar were performed using Fiji software (Schindelin et al., 2012). Basic spectral analysis of the regions of interest (ROIs) in the hyperspectral images of fixed and live cells was carried out using the "Plot Z-axis" tool in Fiji. ROIs were placed in the nucleus and cytoplasm to capture cellular regions with a higher proportion of DNA (nucleus) or RNA (cytoplasm). The color scale for lambda channels was generated using Zeiss Zen Black v. 2.3 software.

#### 1.6.1. Spectral phasor plot analysis and interpretation

For hyperspectral data analysis, we used the spectral phasor transformation using SimFCS software (G-Soft, Illinois-USA) developed at the Laboratory for Fluorescence Dynamics (www.lfd.uci.edu). Through this approach, based on the Fourier transform, data of each pixel of spectral images is processed as described previously by (Malacrida et al., 2016). The spectral phasor transform is defined as:

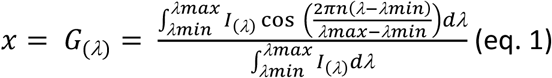

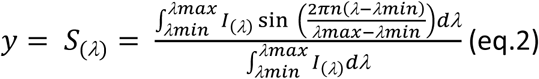

I(λ) is the intensity at each step in a spectral range λ max to λ min, n the harmonic number (1 in our case), λ min is the initial wavelength. Values of G and S are assigned to each pixel in the images and located in the spectral phasor plot. SimFCS software calculates a 2D phasor plot and an average intensity image (from the 22-channel spectral image). In the phasor plot, spectral data is represented as a cloud, whereas the heat map shows the accumulation of pixels. The connection between real and imaginary space can be exploited using cursors in the phasor plot. By doing so, one can interactively understand the fluorescent components presented in the sample.

The position at the phasor plot depends on the spectrum shape, where spectrum center of mass is captured by the phase angle (θ), and the spectrum width by the modulation vector (M). Spectral shift to longer wavelengths implies increasing phase angle and a spectrum broadening moves the position toward the center (M decreases). One of the most significant properties of the phasor plots is the use of the vector properties, such as the linear combination and the reciprocity principle. The linear combination allows for the study of multiple components in a mixture as a sum of fractions of single components. Thus, the position of a combination of two spectra are located on the line that connects the two pure components (Malacrida et al., 2016). The reciprocity principle allows for the selection of a region of interest (ROI, or cursors) on the phasor plot and identifies the corresponding pixels in the average intensity. Additionally, this phasor property also operates in reverse by drawing an ROI on the original image and obtaining the associated phasor plot. We utilized this property to map the AO spectral properties within the cellular environments by masking nuclear or cytoplasmic ROIs. The Phasor plot enables the use of a median filter (3x3 kernel) to improve signal to noise; using SimFCS software we applied twice such filter for all our phasor analyses.

For fixed or live cells spectral images, we defined two or three cursors based on phasor position from pure components. In the case of magenta and green cursors (AO fully bound to RNA or DNA respectively) the diameter selected in the SimFCS software was initially set to 0.03 for reference measurements. For the subsequent analysis of fixed and live cell images, the same cursor positions were maintained, but the cursor diameter was increased to 0.05. This adjustment was necessary to account for differences in the signal-to-noise ratio (SNR) between samples. In live cell acquisitions, higher background and lower photon counts result in broader distributions in the phasor plot. Increasing the cursor size allowed us to integrate all relevant pixels within these broadened clusters, ensuring that the complete spectral population was represented in the final analysis. For fixed cells, a two components analysis was used. To quantify the amount of each component at the image, we calculated the component fraction histograms (Ranjit et al., 2017). We determined the fraction of AO bound to DNA (cursors 1) or RNA, counting the number of pixels in the linear combination between two pure components.

For live cells, we used the three components analysis developed by Ranjit et al. (Ranjit et al., 2019) we quantified the fraction of DNA-bound AO, or the third component (C_3_) and its subcellular distribution using the addition rules of phasor plots. Through the segmentation tool of SimFCS software, we segmented individual cells from images and separated nuclei from cytoplasm. Then, a quantitative analysis was made obtaining the contributions of each AO component through fractional histograms. In this representation the number of pixels at each position in the plot is represented along the line or trajectory DNA-RNA and DNA-C_3_, normalized by the total fraction. Then, we could plot the mean value and standard deviation for each histogram.

### 1.7. Statistical analysis

Three independent biological replicates were performed to validate the results. For statistical analysis, 10 images were acquired per condition. Individual cells stained with AO were manually segmented: 15 live cells and 12 fixed cells were analyzed. Mean curves for fraction histograms are presented as mean ± standard deviation.

For a statistical comparison between different groups, we calculated the center of mass (CM) for the AO fraction histogram bound to DNA or C_3_. The CM was calculated as following:

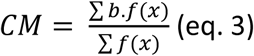

with ‘b’ being the percentage of pixels at the fraction ‘f(x)’ of the variable ‘x’ namely AO- DNA fraction or AO-C_3_ fraction (Malacrida et al., 2016). Center of mass plots show the median value for each condition, along with data dispersion. Violin plots were generated using PlotsOfData software (Postma & Goedhart, 2019).

As the samples did not follow a normal distribution, the Wilcoxon matched-pairs test was applied to analyze the non-parametric data. For the result statistical comparison, a p-value <0.05 was considered statistically significant. Statistical analysis was conducted using GraphPad Software, Inc.

## 2. Results

To evaluate the interaction between AO and nucleic acids, hyperspectral confocal images were acquired from PBS solutions containing AO mixed with either DNA or RNA. By titrating increasing concentrations of nucleic acids while keeping AO constant, spectral phasor (SP) plots were generated. A shift in the phasor’s center of mass was observed up to a saturation point (10 mg/mL for DNA and 2 mg/mL for RNA) beyond which no further changes were detected. Figure 1A shows the centers of mass for each phasor cloud, marked by cyan, green, and magenta cursors. These results confirm that AO undergoes spectral shifts upon binding to nucleic acids, with a blue-shifted emission (lower phase) relative to free AO. Pseudocolor images derived from cursor selection (Figure 1B) display homogeneous signal distributions, suggesting the absence of AO aggregation. Corresponding spectral plots (fluorescence intensity vs. wavelength) revealed emission peaks near 540 nm for both DNA- and RNA-bound AO. Although the emission spectra of free and bound AO are similar, phasor plots demonstrate clear differences in shape and position (Figure 1A, B), highlighting the sensitivity of the spectral phasor approach to subtle changes in the spectrum’s center of mass.

**Figure 1:**
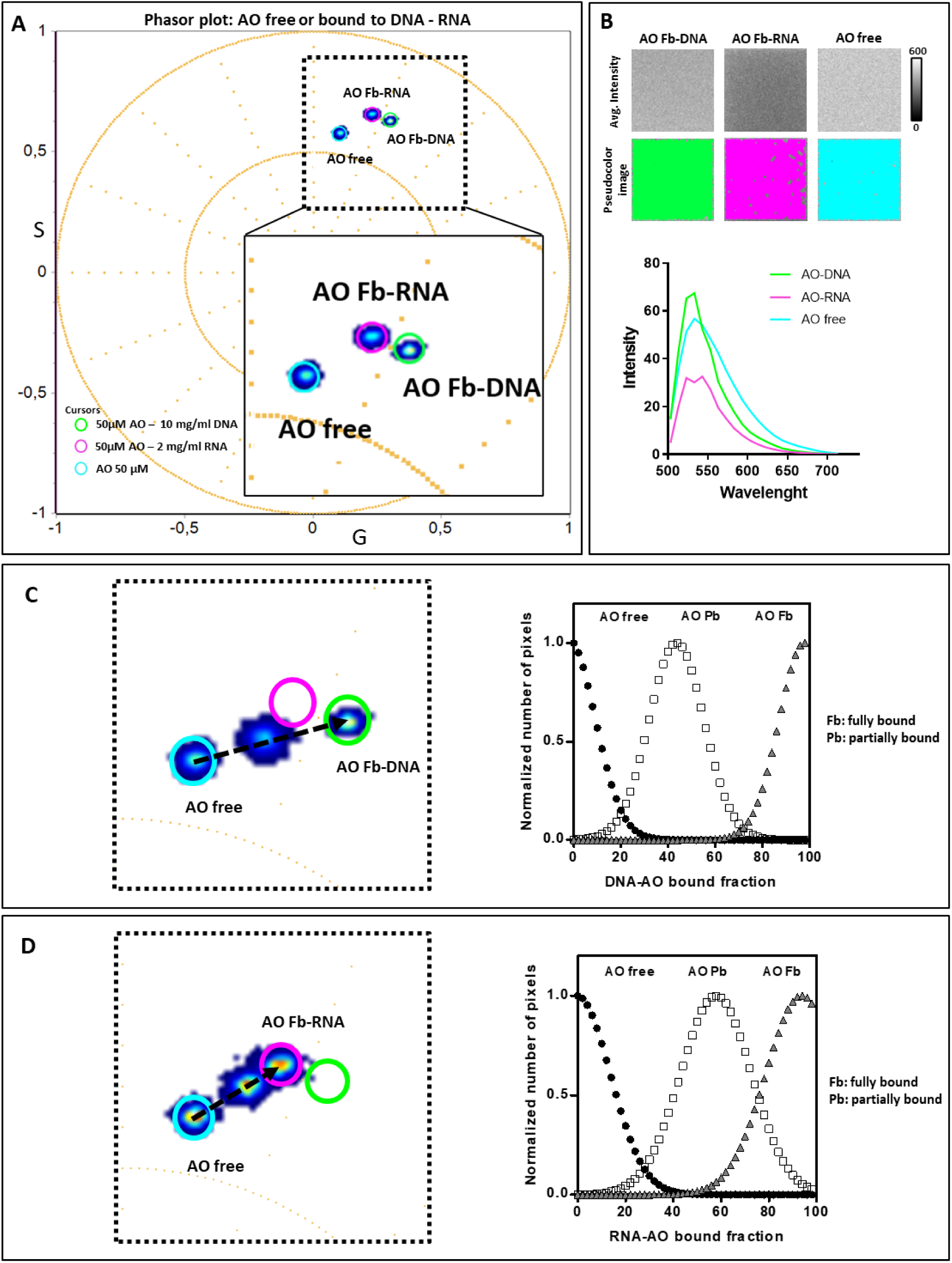
*In vitro* SP characterization of AO interactions with DNA and RNA solutions. (A) SP plot of AO in three solutions, identifying three distinct components. The magenta cursor represents AO fully bound (Fb) to RNA, the green cursor corresponds to AO Fb to DNA, and the cyan cursor indicates free AO. (B) Average intensity and pseudocolor images of the AO solutions shown in (A), illustrating the best fit of the cursor positions to the center of mass of the data distribution. The bottom panel displays the spectral profiles of the images for the three solutions. (C, D) Magnified SP plots highlighting two binding trajectories: partially bound (Pb) and Fb AO with DNA (C) or RNA (D). The corresponding scale bar on the right shows the normalized pixel distribution as a function of the DNA- or RNA-bound fraction of AO. DNA Pb concentration: 0.1 mg/ml; RNA Pb concentration: 0.7 mg/ml.

Figures 1C and D illustrate SP plots for intermediate DNA (0.1 mg/ml) or RNA (0.7 mg/ml) concentrations, representing partial AO binding. These points fall along a linear combination between free AO and fully bound AO (DNA or RNA). Fraction histograms (Figure 1C, D, right panels) quantify the proportions of free AO, partially bound AO, and fully bound AO. This analysis demonstrates the capability of fraction histograms to distinguish and quantify different AO-bound states.

Hyperspectral imaging of AO-stained fixed cells enabled the identification of nuclear and cytoplasmic structures, including nucleoli, based on changes in AO brightness (Figure 2A). The average spectrum from nuclear and cytoplasmic regions of interest (ROIs) has not shown strong differences in AO fluorescence (Figure 2A). However, with Phasor plot analyses we can identify AO interactions with DNA and RNA. Figure 2B displays a cloud overlapping previously identified cursors for AO fully bound to DNA and RNA (Figure 1). The pseudocolor image (Figure 2B) highlights AO-DNA (green pixels) and AO-RNA (magenta pixels) distributions. The nucleoplasm predominantly exhibits AO-DNA binding, while the nucleoli and cytoplasm show higher AO-RNA proportions.

**Figure 2:**
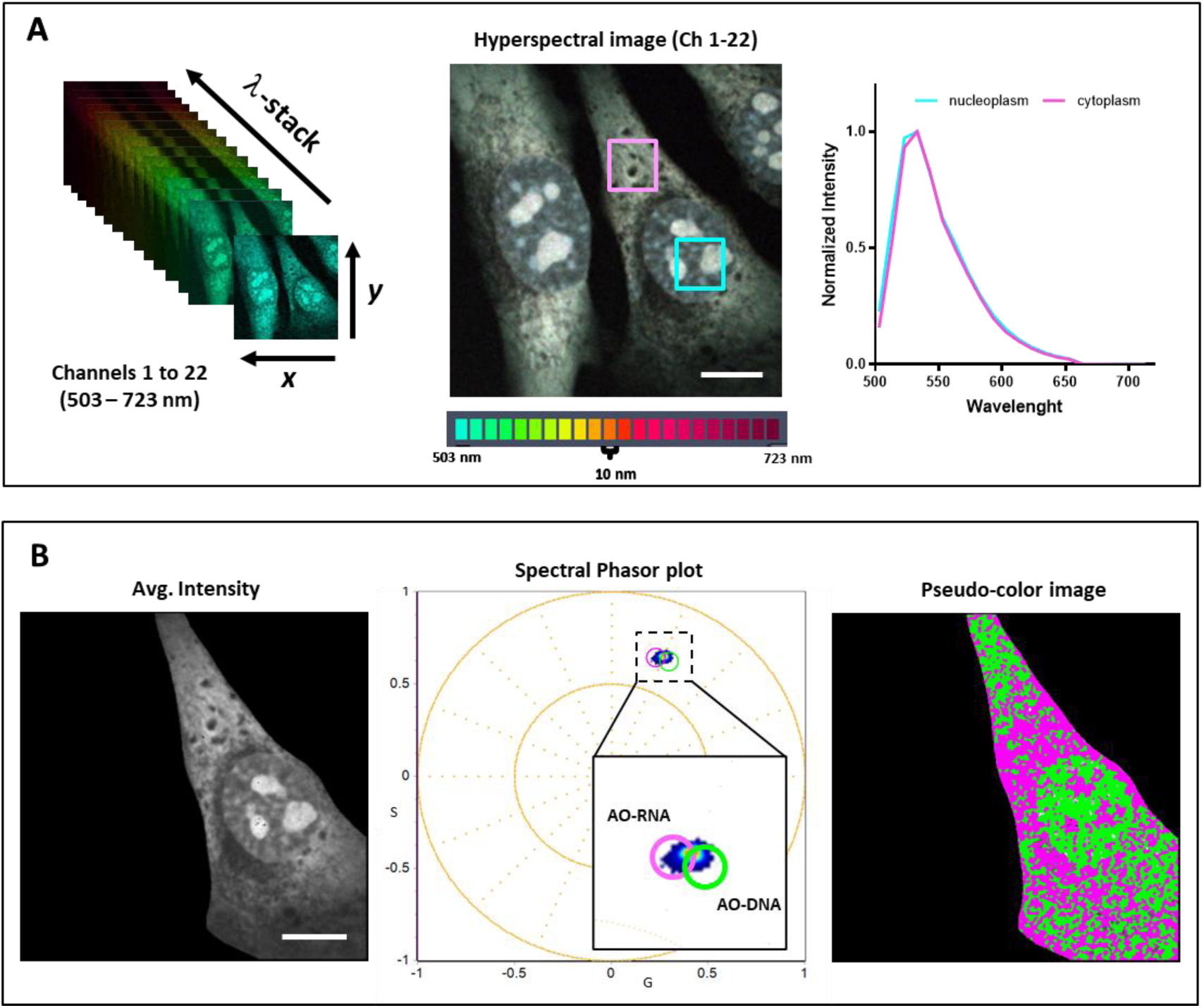
Spectral curves and SP plot analysis of hyperspectral confocal images of AO-stained NIH-3T3 fixed cells. (A) Spectral acquisition across 22 channels (503–723 nm) using confocal microscopy. The hyperspectral image was generated by summing all channels. ROIs were placed on the cytoplasm (magenta) and nucleoplasm (cyan). The graph shows normalized spectral curves for each ROI. (B) Left: Average intensity image of a cell segmented from (A). Middle: Spectral phasor plot. Right: Pseudocolor image showing RNA-rich (magenta) and DNA-rich (green) regions based on phasor cursor positions defined from Figure 1. Cursor diameters were increased to include all pixels, keeping center positions unchanged. Scale bar: 10 µm.

Quantitative SP analysis of AO fluorescence in fixed cells is shown in Figure 3. A pseudocolor image (Figure 3A) indicates DNA-bound AO in the nucleus (red) and RNA-bound AO in nucleoli (greenish/blueish pixels). Cytoplasmic regions predominantly exhibit RNA-bound AO. Fraction histograms of AO-DNA binding (Figure 3B) indicate that, on average, 50% of pixels correspond to equal AO-DNA and AO-RNA proportions across ten analyzed hyperspectral images. Masking analyses of nuclei and cytoplasm (Figures 3C, D) reveal distinct binding patterns. Nucleus masking (Figure 3C) shows a higher fraction of AO-DNA (65% AO-DNA to 35% AO-RNA), confirmed by SP and histogram analyses. Cytoplasmic masking (Figure 3D) shows increased AO-RNA proportions (40% AO-DNA to 60% AO-RNA), visualized as blue pixels in pseudocolor images. Violin plots (Figure 3E) confirm significant differences in DNA-bound AO proportions between the nucleus and cytoplasm (n=10, p=0.0005).

**Figure 3:**
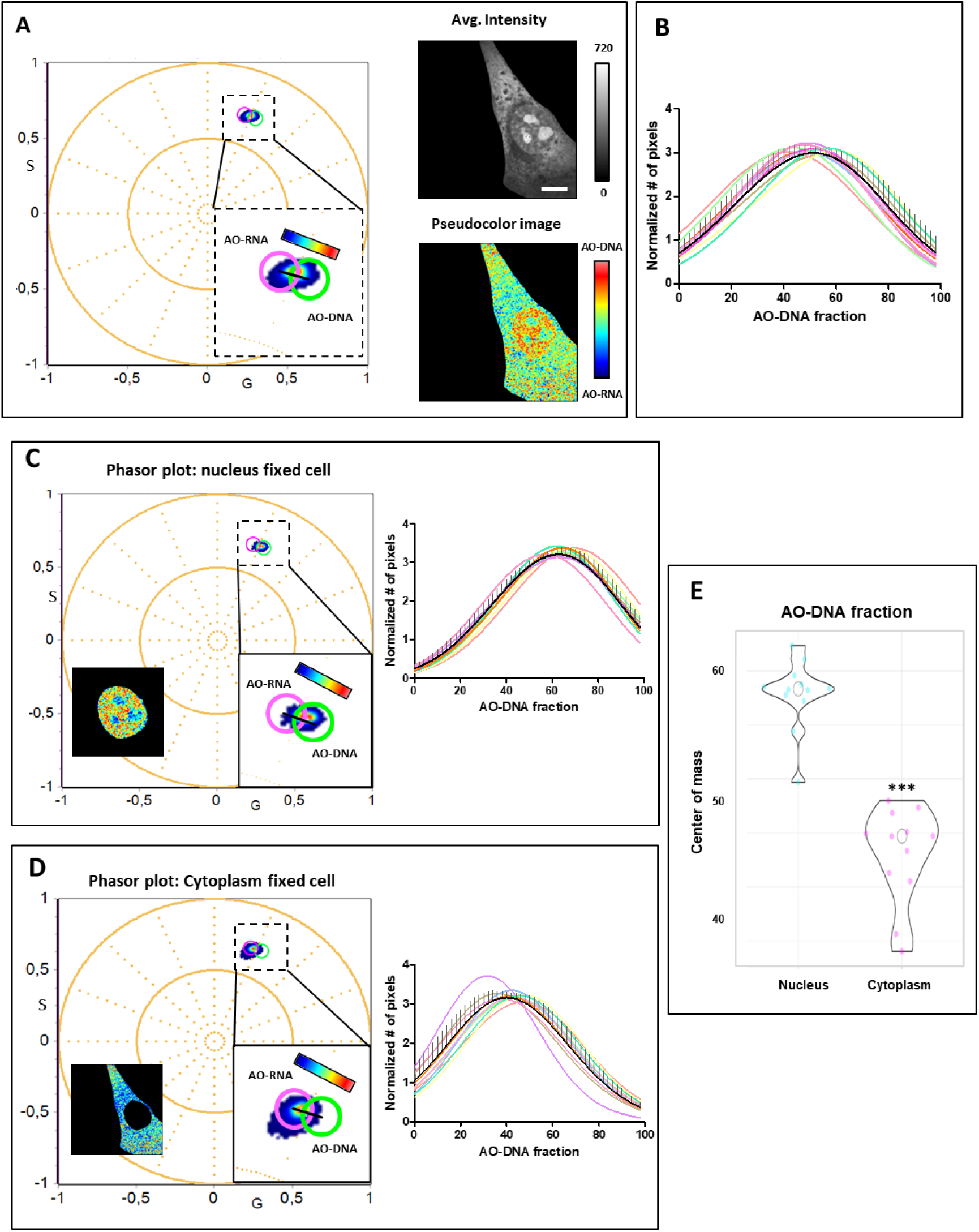
Two-component SP plot analysis of AO-stained fixed NIH-3T3 cells. (A) SP of the same cell analyzed in Fig. 2. SP showing AO-DNA and AO-RNA distribution along the black line trajectory. Pseudocolor image and color scale bar (right) indicate pixel proportions, from red (DNA-rich) to blue (RNA-rich). (B) Histograms showing the normalized number of pixels as a function of the AO-DNA fraction from 10 cells (colored curves), with mean (black) ± SD, showing ∼50% DNA-binding on average. (C, D) Phasor analysis of nucleus (C) and cytoplasm (D) from the SP analysis of segmented nucleus (C) and cytoplasm (D) from the same representative cell in (A). Pseudocolor images and histograms depict the spatial distribution of AO-DNA and AO-RNA. (E) Violin plots of AO-DNA center of mass in nucleus and cytoplasm. Wilcoxon matched-pairs test: ***p = 0.0005, N = 10. Scale bar: 10 µm.

In live cells, AO fluorescence reveals nuclear structures with green fluorescence, including well-defined nucleoli, and weak green cytoplasmic fluorescence, except for the granules with intense orange to red fluorescent (Figure 4A). Spectral plots (Figure 4A, right) show a second emission peak at 640 nm, indicative of cytoplasmic AO particulates. Dividing hyperspectral images into two sub-images (Figure 4B) highlights nuclear signals (503-583 nm) and cytoplasmic particulates (603-723 nm). SP analysis (Figure 4C) identifies three components: AO-DNA (green), AO-RNA (magenta), and a third component (C_3_, cyan rectangle). The pseudocolor image (Figure 4C, right) shows AO-DNA predominantly in the nucleus, AO-RNA in nucleoli and cytoplasm, and C_3_ in cytoplasmic regions.

**Figure 4:**
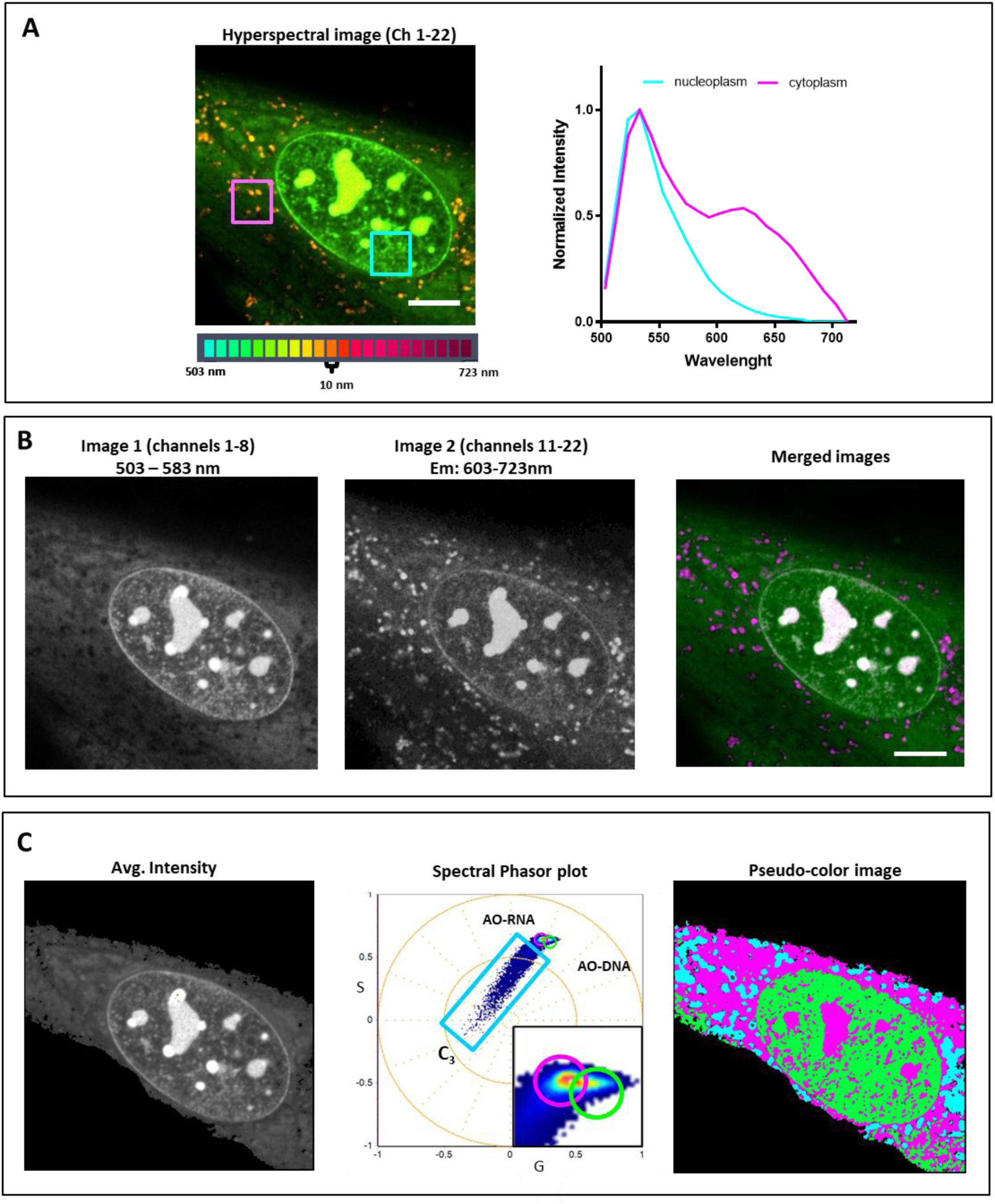
Spectral and phasor analysis of AO-stained live NIH-3T3 cells. (A) Hyperspectral confocal image (sum of 22 channels) with ROIs on nucleoplasm (cyan) and cytoplasm (magenta); right: average AO spectra for each ROI. (B) Left and center: average intensity projections from two halves of the lambda stack. Right: merged images (green = image 1, magenta = image 2). (C) Center: SP revealing three components, AO-RNA (magenta), AO-DNA (green), and a third component (C_3_). Right: pseudocolor image showing C_3_-related pixels in cyan. Left: Average intensity image of the lambda stack. Scale bar: 10 µm.

By combining phasor plots with three-component analysis, the DNA/RNA AO fractions and C_3_ distribution were resolved. AO-DNA binding is highest in the nucleus, while RNA-bound AO is prominent in the cytoplasm (Figure 5A). Intermediate proportions are observed in nucleoli. Fraction histograms of DNA-bound AO in live cells (Figure 5B) show a higher average fraction (80-90%). Masking the nucleus reveals strong variability in AO binding fractions (Figure 5C). The cytoplasmic region shows a distinct C_3_ presence (Figure 5D). Violin plots (Figure 5E) confirm that C_3_ is significantly more abundant in the cytoplasm than the nucleus (median 12%, p<0.0001).

**Figure 5:**
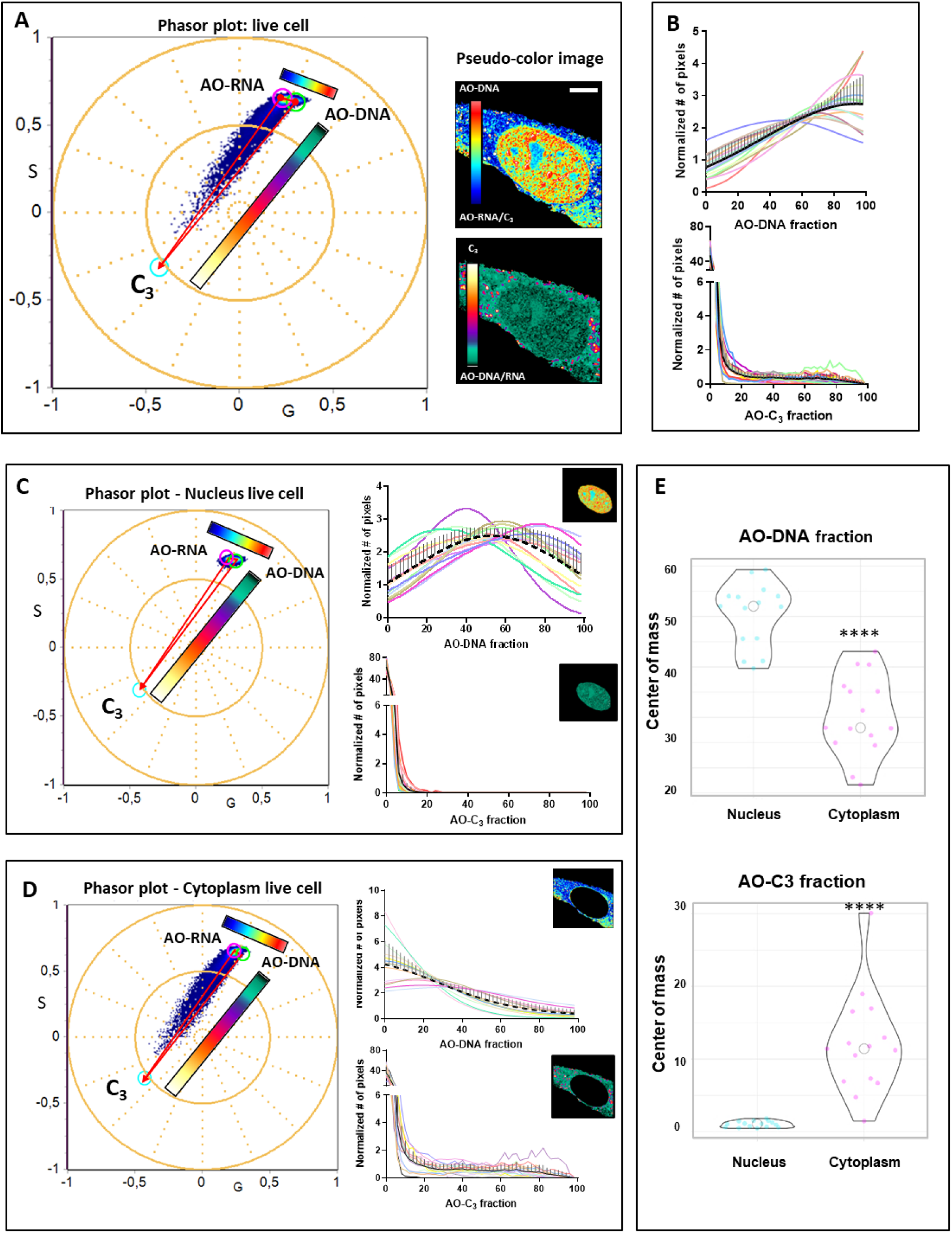
Three-component SP analysis of AO-stained live NIH-3T3 cells. (A) Phasor plot of a representative cell showing two projections: AO-DNA vs. AO-RNA, and AO-DNA vs. a third component (C_3_). Pseudocolor images (right) illustrate spatial distribution and proportion of components in each dimension. (B) Histograms of pixel fractions (%) for AO-DNA and C3 across 15 cells; mean ± SD in black. (C, D) Three-component phasor analysis of segmented nuclei (C) and cytoplasm (D), with corresponding pseudocolor images from (A). Histograms show component distributions. (E) Violin plots of AO-DNA and C_3_ center of mass in nucleus vs. cytoplasm. Wilcoxon matched-pairs test: ****p < 0.0001, N = 15. Scale bar: 10 µm.

To explore C_3_’s nature, live cells were co-stained with AO and LysoTracker Deep Red (Figure 6A-E). Although AO marks acidic compartments, not all AO-positive structures correspond to acidic organelles. LysoTracker appears in the fourth SP quadrant, enabling linear combinations with AO components (Figure 6F). Co-stained cells (Figure 6G) show complex SP distributions, with distinct clusters for C_3_ and combinations of LysoTracker with RNA/DNA-bound AO. Pseudocolor images reveal two subcellular populations: AO-labeled acidic organelles and RNA/DNA-bound structures. These results emphasize the specificity of AO for different subcellular components and its ability to distinguish non-acidic cytoplasmic structures.

**Figure 6:**
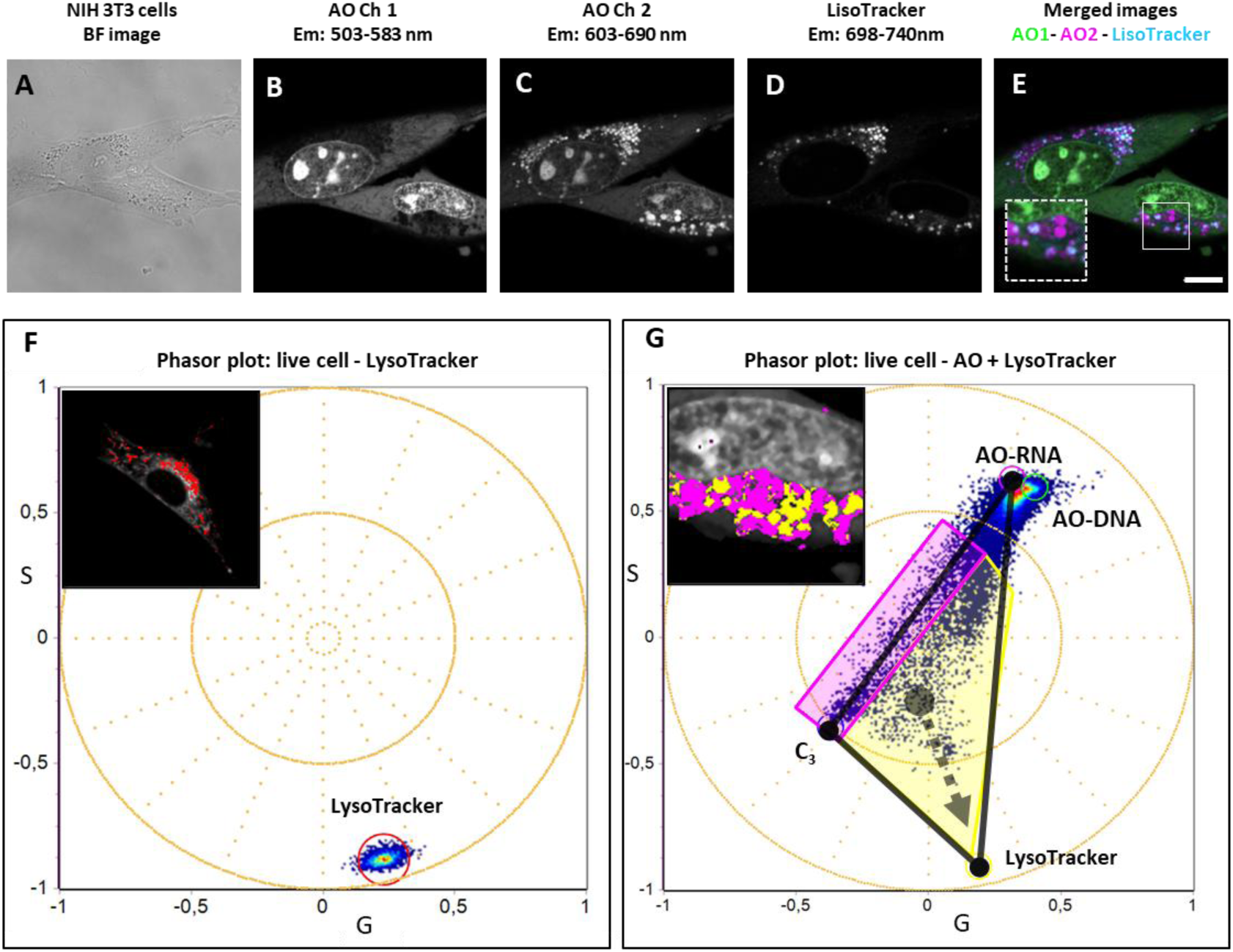
Characterization of the third AO component (C_3_) in live NIH-3T3 cells. (A) Brightfield image. (B-D) Maximum intensity projections of spectral confocal images of cells stained with AO and LysoTracker Deep Red (Ex: 647 nm). The original lambda stack was separated into three channels: (B) Channel 1 (AO), (C) Channel 2 (AO), and (D) Channel 3 (LysoTracker). (E) Merged image: green (AO Ch1), magenta (AO Ch2), cyan (LysoTracker). Zoomed inset shows cyan-magenta spots, acidic compartments co-stained with AO and LysoTracker. (F) Phasor plot of a cell stained with LysoTracker only. Red cursor marks the lysosome-like cluster (see inset image). (G) Phasor plot of a cell stained with both AO and LysoTracker. Yellow region highlights pixels combining AO-RNA, AO-DNA, and C_3_ with LysoTracker. Magenta region indicates AO-RNA and C_3_ without LysoTracker. Inset image links phasor regions to magenta and cyan zones in (E). Scale bar: 10 µm.

## 3. Discussion

The interaction of AO with nucleic acids *in vitro* has been extensively studied and characterized through spectroscopic and molecular analyses (Amado et al., 2017; Plemel et al., 2017; Sayed et al., 2016). The spectral changes associated with AO’s interaction with itself or with nucleic acids (DNA and RNA) provide a valuable tool for the qualitative and semi-quantitative differentiation of sub-cellular structures in living and fixed cells. In this study, we revisited AO fluorescence in cells using the unique, model-free SP analysis of hyperspectral imaging. This approach proved sensitive to diverse AO environments within cells, enabling the identification of multiple sub-cellular compartments and providing a comprehensive understanding of AO’s molecular interactions within cellular contexts.

Despite similarities in the spectral shifts experienced by AO when bound to RNA or DNA, both in terms of spectrum maximum and shape, SP analysis offers a clear distinction in the phasor plot due to the specific molecular environments provided by RNA or DNA. Specifically, AO interactions with RNA generate a spectral shift towards longer phases (redder spectra) compared to DNA. These spectral differences are attributed to AO’s diverse modes of interaction with nucleic acids, such as single-strand (ss) or double-strand (ds) binding, base-pair intercalation, monomeric binding, and stacking interactions leading to dimer formation (Amado et al., 2017; Delic et al., 1991; Kapuscinski & Darzynkiewicz, 1987). To refine our understanding of these interactions, we conducted *in vitro* studies with AO solutions and purified ss/ds DNA and RNA. These experiments revealed the unique SP fingerprints corresponding to distinct molecular environments.

Solutions of AO mixed with varying concentrations of nucleic acids were analyzed using SP. Results shown in Figure 1 confirmed that solutions comprising free and bound AO align along a line connecting pure components, underscoring the linear combination principle inherent in phasor plots. The overlap of phasor positions between *in vitro* and cellular environments further validates the robustness of this model-free analysis, enabling direct comparisons of AO’s spectroscopic properties across experimental systems.

Plemel et al. reported that a 140 ng/µl RNA solution combined with 50 µM AO formed AO-RNA complexes exhibiting a secondary peak at approximately 640 nm (Plemel et al., 2017). However, our experiments, even at higher RNA concentrations (2 µg/µl), did not produce this spectral peak. This discrepancy may stem from the different excitation wavelengths used. We selected 488 nm excitation, commonly employed in studies involving AO-stained living and fixed cells (Fucic et al., 2021; Kasten,1999; Plemel et al., 2017; Zelenin, 1999), whereas Plemel et al. used 457 nm excitation for *in vitro* solutions. The 457 nm excitation preferentially excites dimeric AO forms and B-type complexes associated with single-stranded RNA, which may explain the observed spectral differences (Zelenin, 1999). Other factors, such as extended incubation times or varying dye-to-solute ratios, could also influence AO interactions. For instance, Kapuscinski et al. noted that AO’s interaction with RNA can form precipitates fluorescing at 650 nm over prolonged incubation (Kapuscinski et al., 1982).

Fixed NIH-3T3 cells stained with AO exhibited a homogeneous spectral profile with only subtle differences between nuclear and cytoplasmic regions (Figure 2A). This observation contrasts with previous reports, where AO staining in fixed cells produced a pattern of green nuclei, yellow nucleoli, and red cytoplasm (Kasten,1999; Plemel et al., 2017; Zelenin, 1999). The use of Triton X-100 post-fixation in our protocol likely contributed to the loss of the red spectral component. Nevertheless, our SP analysis effectively differentiated RNA- and DNA-bound AO molecular environments under these conditions (Figure 2B).

The SP analysis of AO hyperspectral images in live cells further highlights the versatility of this approach. Without assuming a priori models, we identified a third molecular environment (C_3_) for AO within the phasor space. The variability among AO-DNA and AO-RNA, components within a population of cell nuclei may reflect differences in cell cycle stages, as our cultures were unsynchronized. This method could provide valuable insights into cellular processes, such as apoptosis, previously explored using AO in fixed-cell experiments (Plemel et al., 2017).

Based on prior studies, AO’s interaction with acidic compartments in living cells induces a metachromatic shift (Delic et al., 1991; Kasten,1999; Plemel et al., 2017; Thomé et al., 2016; Zelenin, 1999). In our results, AO and LysoTracker Deep Red co-labeled live cells revealed that some C_3_-positive pixels correspond to acidic organelles, while others represent AO-stained organelles containing RNA. These organelles likely encompass structures such as stress granules, processing bodies, ribosomes, and RNA-associated extracellular vesicles like exosomes (Protter & Parker, 2016; Riggs et al., 2020).

In summary, SP analysis of hyperspectral imaging offers a powerful, model-free method to study AO’s interactions with nucleic acids and sub-cellular structures. By providing unique molecular fingerprints for AO in diverse environments, this approach enhances our understanding of AO’s binding dynamics and its application in cell biology.

## 4. Conclusion

Overall, SP analysis of AO hyperspectral data has proven to be an exceptionally sensitive and effective tool for exploring the intricate photophysics of AO within various sub-cellular compartments, offering greater sensitivity and insight than analyses based solely on spectral information. This method enabled the quantitative analysis of DNA- and RNA-bound AO fractions and highlighted distinct differences in SP profiles between fixed and live cells. In live cells, a third molecular environment was identified, associated with the 640 nm AO peak, likely corresponding to a subset of organelles. However, the precise identity of some of these organelles remains unclear, presenting avenues for future research.

SP analysis surpasses classical methods of radiometric or emission spectral evaluation, providing a robust capability to distinguish acidic compartments, where AO forms multimers, from cytoplasmic structures interacting with RNA. By integrating AO staining with the SP approach, whether in fixed or live samples, researchers gain access to a powerful and efficient tool for analyzing complex sub-cellular organization across diverse biological systems, ranging from individual cells to tissues.

## Acknowledgments

The authors declare no conflict of interests. The authors would like to thank the Advanced Bioimaging Unit-Institut Pasteur of Montevideo-HC Udelar. LM is supported by the grants 2020-225439, 2021-240122, 2022-252604 of Chan Zuckerberg Initiative DAF, an advised fund of Silicon Valley Community Foundation. LM, MD and FRZ were supported by FOCEM - Fondo para la Convergencia Estructural del Mercosur (COF 03/11). FRZ and LM are investigators of PEDECIBA.

## Data availability statement

The data that support the findings of this study are openly available at the following URL/DOI: https://doi.org/10.5281/zenodo.15500562

